# Natural variation identifies new effectors of water use efficiency in *Arabidopsis*

**DOI:** 10.1101/2022.03.28.486113

**Authors:** Govinal Badiger Bhaskara, Jesse R Lasky, Samsad Razzaque, Li Zhang, Taslima Haque, Jason E Bonnette, Guzide Zeynep Civelek, Paul E Verslues, Thomas E Juenger

**Author notes:** **Corresponding authors:** Govinal Badiger Bhaskara, Thomas E Juenger. **Email:**.

## Abstract

Water use efficiency (WUE) is the ratio of biomass gained per unit of water consumed; thus, it can be altered by genetic factors that affect either side of the ratio. In the present study, we exploited natural variation for WUE as an unbiased approach to discover loci affecting either biomass accumulation or water use as factors affecting WUE. Genome-wide association (GWAS) analysis using integrated WUE measured through carbon isotope discrimination (δ^13^C) of *Arabidopsis thaliana* accessions identified genomic regions associated with WUE. Reverse genetic analysis of 70 candidate genes selected based on the GWAS results and transcriptome data identified 25 genes affecting WUE as measured by gravimetric and δ^13^C analyses. Mutants of four genes had higher WUE than wild type, while mutants of the other 21 genes had lower WUE. The differences in WUE were caused by either altered biomass or water consumption (or both). Stomatal density was not a primary cause of altered WUE in these mutants. Leaf surface temperatures indicated that transpiration differed for mutants of 16 genes, but generally biomass accumulation had greater effect on WUE. The genes we identified are involved in diverse cellular processes including hormone and calcium signaling, meristematic activity, photosynthesis, flowering time, leaf/vasculature development, and cell wall composition; however, none of them had been previously linked to WUE or traits related to plant water relations. Thus, our study successfully identified new effectors of WUE that can be used to understand the genetic basis of WUE and improve crop productivity.

## Introduction

Plants move large amounts of water through the soil-plant-atmosphere continuum from water uptake by roots to loss of water to the atmosphere through stomatal pores. Plants can limit water loss by closing stomata or by having fewer or smaller stomata. However, this must be balanced with the need to maintain sufficient gas exchange for CO_2_ to enter the leaf for photosynthesis (1). If gas exchange through the stomata is too restricted, depletion of internal leaf CO_2_ can limit photosynthesis and hence, plant biomass and productivity (2). Globally, agriculture accounts for over 80% of freshwater use (3). Therefore, improving WUE or enhancing water productivity (i.e., more yield per unit of water consumed) is important for crop improvement (4). However, efforts to improve WUE have met with limited success thus far (5).

Molecular approaches have demonstrated that reduced stomatal conductance (*g*_*s*_) can improve WUE without reducing biomass accumulation (6, 7). Since abscisic acid (ABA) is key inducer of stomatal closure, several studies have targeted ABA signaling genes to reduce *g*_*s*_. Several transgenic strategies to alter ABA sensitivity led to increased WUE while maintaining high biomass and grain yield under progressive drought (8–10). Ectopic expression or mutation of genes which regulate stomatal development can also lead to increased WUE via effects on stomatal density (6). While these studies successfully increased WUE by manipulating previously described regulatory genes, other approaches are needed to allow an unbiased and broader search for new effectors of WUE. Particularly, the studies mentioned above focused on manipulating the transpiration side of WUE by changing stomatal density or altering stomatal opening and closing behavior. It is less clear whether changing the other side of WUE, biomass accumulation, can also be a path to increased WUE. Because of the complexity of plant growth regulation, this side of the WUE is less amenable to manipulation by targeting *a priori* candidate genes.

Plants exhibit natural genetic variation in WUE (11–17). This includes both variation in the transpiration (stomatal behavior and density) as well as the biomass (biomass and plasticity of biomass during water limitation) sides of WUE (7). Quantitative trait loci (QTL) mapping has identified many QTL driving variation in δ^13^C (the ratio of ^13^C to ^12^C), a widely used proxy for integrated WUE (7). One such study found that a single amino acid change in *Mitogen-Activated Protein Kinase 12 (MPK12)* is responsible for variation in stomatal conductance and WUE in a number of Arabidopsis accessions (18, 19). Another QTL analysis discovered that *ERECTA*, a gene involved in stomatal development, contributes to variation in δ^13^C and transpiration efficiency (20). Genome wide association (GWAS) using large panels of natural accession is an attractive alternative to bi-parental linkage mapping. In Arabidopsis, there is substantial natural variation in many water use and drought-related traits (21–24) and the decay of linkage disequilibrium occurs over a relatively short interval. Thus, once SNPs associated with a particular trait are identified, testing only a few candidate genes in the vicinity of strongly associated SNPs, or cluster of moderately associated SNPs, can find gene(s) affecting the trait of interest (23, 24). Integrating other information, such as transcriptome data, can further assist in identifying the most promising candidate genes (25). The combination of GWAS and reverse genetic testing of candidate genes allows for an open-ended search for effector genes. This is particularly useful for traits, such as WUE, where it is difficult to apply traditional forward genetic screening. To date, only a few studies have employed GWAS to identify candidate loci affecting WUE (25–29) and there is relatively little data on validation of candidate genes identified by GWAS for the drought-related traits (23, 24).

We applied GWAS and reverse genetic testing to discover loci affecting WUE plasticity whereby Arabidopsis accessions increase, or fail to increase, WUE in response to soil drying. Such an approach allowed us to find genes that had not been previously implicated in WUE including genes affecting the biomass accumulation side of WUE rather than the transpiration side. Indeed, the largest effect we observed was in mutants of *Plant Cysteine Oxidase 5 (PCO5)*, which affects WUE mainly by allowing increased biomass production. Our study identified several new WUE effector genes that offer new routes to manipulate and improve this key trait

## Results

### GWAS Analysis to Identify Genomic Regions Associated with Variation in WUE Plasticity

We leveraged the measurements of integrated WUE as δ^13^C of above ground biomass of well-watered plants or plants subjected to terminal drought from Kenney et al. (22). We reanalyzed the core subset of 185 accessions (dataset S1). Most accessions had a higher WUE in the drought treatment (higher δ^13^C, F = 42.89, p < 0.0001; Fig. 1A). Only a few accessions had decreased or unchanged WUE in the drought treatment compared to the unstressed control (Fig. 1A). GWAS was performed using the plasticity of WUE (Fig.1B; Plasticity = drought –well-watered control; mean of −0.585, and range −3.35 to 1.75) as well as the WUE of the well-watered control and drought treatments (Fig. S1). While all three sets of GWAS data identified potentially interesting associations, we focused on the WUE plasticity GWAS for further study since the ability to change WUE in response to changing environmental conditions is thought to be an important adaptive trait whose regulation is poorly understood. The SNPs most significantly associated with WUE plasticity were distributed across several genomic regions (Fig. 1C). As GWAS generates many candidates, selecting the most promising genes for the follow-up functional study is challenging. Thus, we used several layers of data to identify and prioritize candidate genes (Fig. S2A). We first generated a list of candidate genes for which any part of the gene body (UTRs, introns and exons) was within 5 kb of one or more of the top 500 lowest-*P-* value SNPs (nominal *P* = 4.37 × 10^−5^ to 4.2 × 10^−3^). The 1058 genes so identified (Dataset S2) were then categorized based on their stress-induced gene expression to further determine which genes were most likely to be involved in WUE plasticity. Stress-responsive changes in gene expression were identified by referring to transcriptomic studies (30, 31, Dataset S3 and S4) that examined gene expression during acclimation to an extended duration of non-lethal drought stress conditions similar to those used to measure WUE plasticity in the current study. This analysis identified 198 genes that were associated with the top 500 WUE plasticity SNPs and that were upregulated during drought stress (Fig. S2B, Dataset S5). These genes were further subjected to Gene Ontology (GO) enrichment analysis which identified 10 significant functional annotation terms (Fig. S2C and S3, Dataset S6). To incorporate a range of gene functions into our validation experiments, we selected the top 10 genes from each enriched GO term based on lowest-*P-* values of their associated SNPs. We also included all 19 genes from the transcriptional regulation GO cluster. This filtering identified 72 candidate genes (Dataset S7). One limitation of this approach is that it may discard poorly annotated genes. We found 29 such genes annotated as proteins of unknown functions associated with the top 500 SNPs and were transcriptionally responsive to stress (Dataset 5: genes highlighted in red). Thus, we also included those 29 genes in our functional study.

**Figure. 1.**
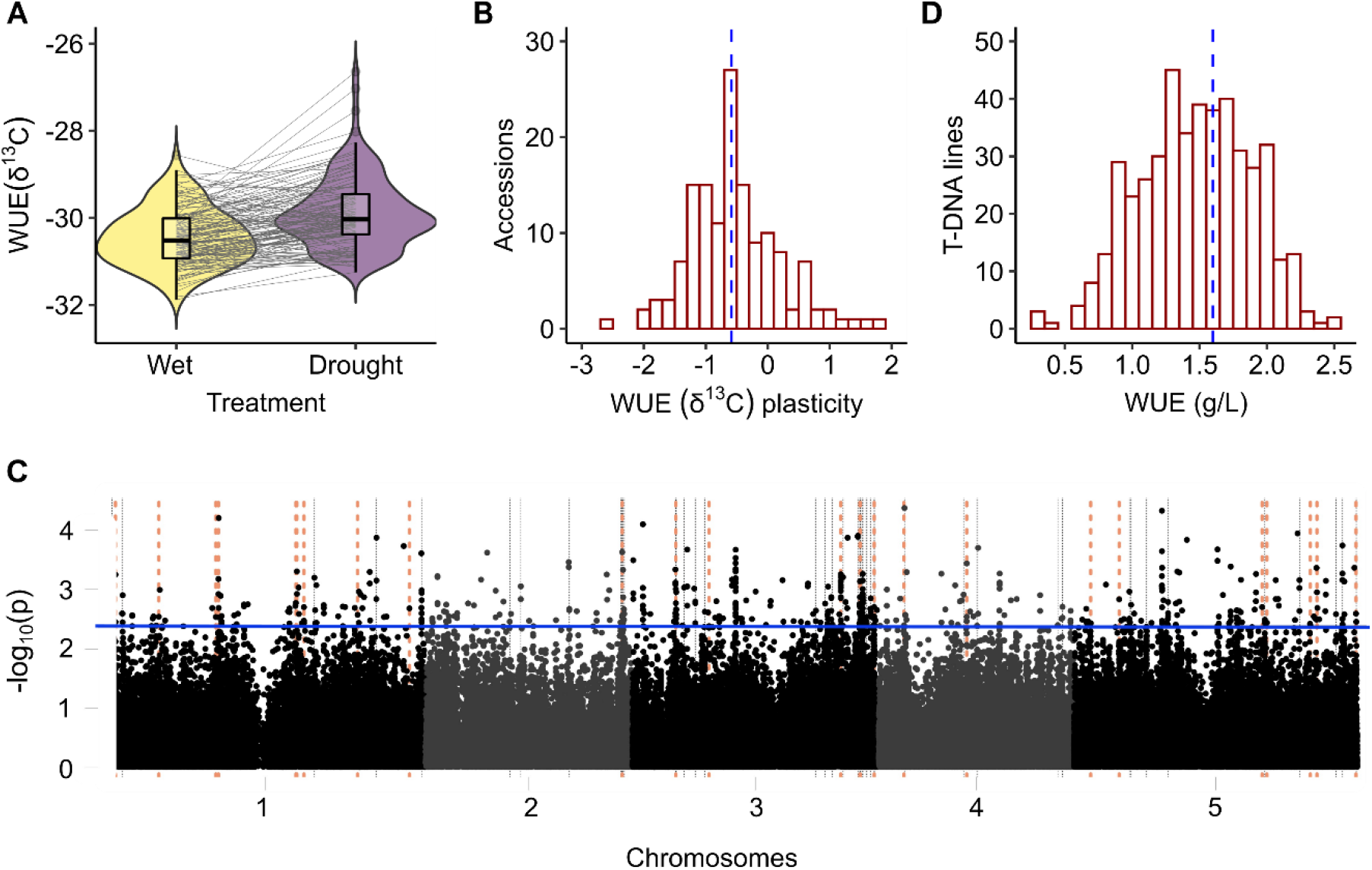
Natural variation, GWAS analysis, and reverse genetic tests of candidate genes for WUE by gravimetric and δ^13^C analyses. (*A*) Plastic response of 185 *Arabidopsis* accessions to terminal drought. WUE (δ^13^C: ^13^C isotopic composition of above ground biomass) values are pooled accession values within each treatment. (*B*) Distribution of WUE (δ^13^C) plasticity (difference in δ^13^C between drought and wet) for 180 *Arabidopsis* accessions. Blue vertical line indicates the median. (*C*) Manhattan plot of SNP *P* values from the GWAS analysis using the WUE plasticity data shown in *A*. The blue horizontal line indicates the cutoff for the top 500 low-*P*-value SNPs. The vertical lines indicate the genomic positions for the prioritized GWAS candidates for which we analyzed WUE (biomass produced per unit of water consumed) using reverse genetics. Grey lines indicate location of genes where T-DNA mutant(s) did not have significantly altered WUE and orange dotted lines indicate genes where the T-DNA mutant(s) did have significant effect on WUE (*D*) Distribution of gravimetric WUE (ratio of biomass to water consumed) for 88 T-DNA mutants from high throughput screening using small containers (*n* = 3-5 biological replicates per genotype). Blue vertical line indicates the mean of Col −0 wild type.

### Reverse Genetic Analysis of Prioritized Genomic Regions

We isolated 88 homozygous T-DNA insertion lines covering 70 candidate genes (we were unable to obtain homozygous T-DNA lines for the other 31 prioritized genes. See methods for the details). These homozygous lines were tested for their effect on WUE by gravimetric analysis using a high throughput system (Fig. 1D and B, Fig. S4A and B). This first round of screening identified mutants for 27 genes that significantly differed from wild type for WUE (Fig. S4A, Dataset S9 and S10). Among these 27 genes, only 9 genes had previously known roles in abiotic stress (Dataset S11).

These mutants were further analyzed for their effect on WUE by two independent gravimetric analyses using larger containers (237 mL) to allow the plants a larger rooting volume and space for rosette biomass (Fig. S4C). We also measured δ^13^C of aboveground biomass, leaf surface temperatures, stomatal density (SD) and stomatal index (SI). This set of measurements included *mitochondrial editing factor11* (*mef11-5*) and *open stomata kinase* (*ost1-3;* OST1 is also referred to as SnRK2.6) mutants as controls known to affect water loss. *mef11-5* is hypersensitive to ABA-mediated stomatal closure and exhibits elevated leaf temperatures and reduced water loss under water stress (32, 33). In contrast, *ost1-3* has reduced ABA sensitivity leading to reduced stomatal closure, greater leaf water loss, and reduced leaf temperature (34, 35). Note that mutants for two genes, *Cytochrome P 450 family 76* (*CYP76C2*) and *AT3G58660* did not germinate for unknown reasons, and so we were unable to include them in our further study. We found a significant positive correlation (*R*^*2*^= 0.627, *P <* 0.001) for WUE between smaller containers (high throughput system) and larger containers for these mutants (Fig. S5A).

These experiments identified genes that altered either side (biomass accumulation or transpiration) of WUE (Fig.2A). For example, *plant cysteine oxidase 5* (*pco5-1, pco5-2*) had a strongly significant (adjusted *P* < 0.05) increase in biomass which gave it a higher WUE despite the fact that it consumed substantially more water than the Col-0 wild type (Fig.2A, purple circles). A few other mutants had no change in biomass but showed moderately (significant based on nominal *P* value) reduced water consumption, leading to increased WUE (Fig.2A, blue circles). Several other mutants had moderately decreased WUE mainly due to decreased biomass accumulation with no significant change in water consumption (Fig. 2, green circles). We also found mutants with strong reduction in biomass as well as water consumption. However, changes in biomass had a relatively larger impact on WUE in these mutants (Fig. 2A, orange circles). The two control mutants *mef11-5* and *ost1-3* showed high and low WUE respectively (Fig. 2A), consistent with their effects on ABA sensitivity. Interestingly, *mef11-5* was the only mutant in our analysis where decreased biomass was associated with increased WUE. Across these mutants, there was a strong correlation between above ground biomass gain and water consumption (Fig. 2B; *R*^*2*^ = 0.89, *P <* 0.001). We also found that WUE was moderately correlated with biomass (Fig. 2C, *R*^*2*^ = 0.628, *P <* 0.001) and weakly correlated with water consumption (Fig. 2D, *R*^*2*^ = 0.364, *P <* 0.001), suggesting that the majority of mutants analyzed primarily altered the biomass side of the WUE ratio. In addition, we found a relatively weak correlation between WUE and stomatal density (SD, Fig. 2E, *R*^*2*^ = 0.142, *P* < 0.017), and no significant relationship between WUE and stomatal index (SI, Fig. S5B, *R*^*2*^ = 0.083, *P <* 0.057) suggesting that changes in SD or SI were not the main driver of altered WUE in our set of mutants. The correlation between gravimetrically determined WUE and δ^13^C was significant and positive (Fig. 2F; *R*^*2*^ = 0.546, *P <* 0.001). We discuss sets of discovered genes based on their impact on WUE, biomass, and water consumption below.

**Figure. 2.**
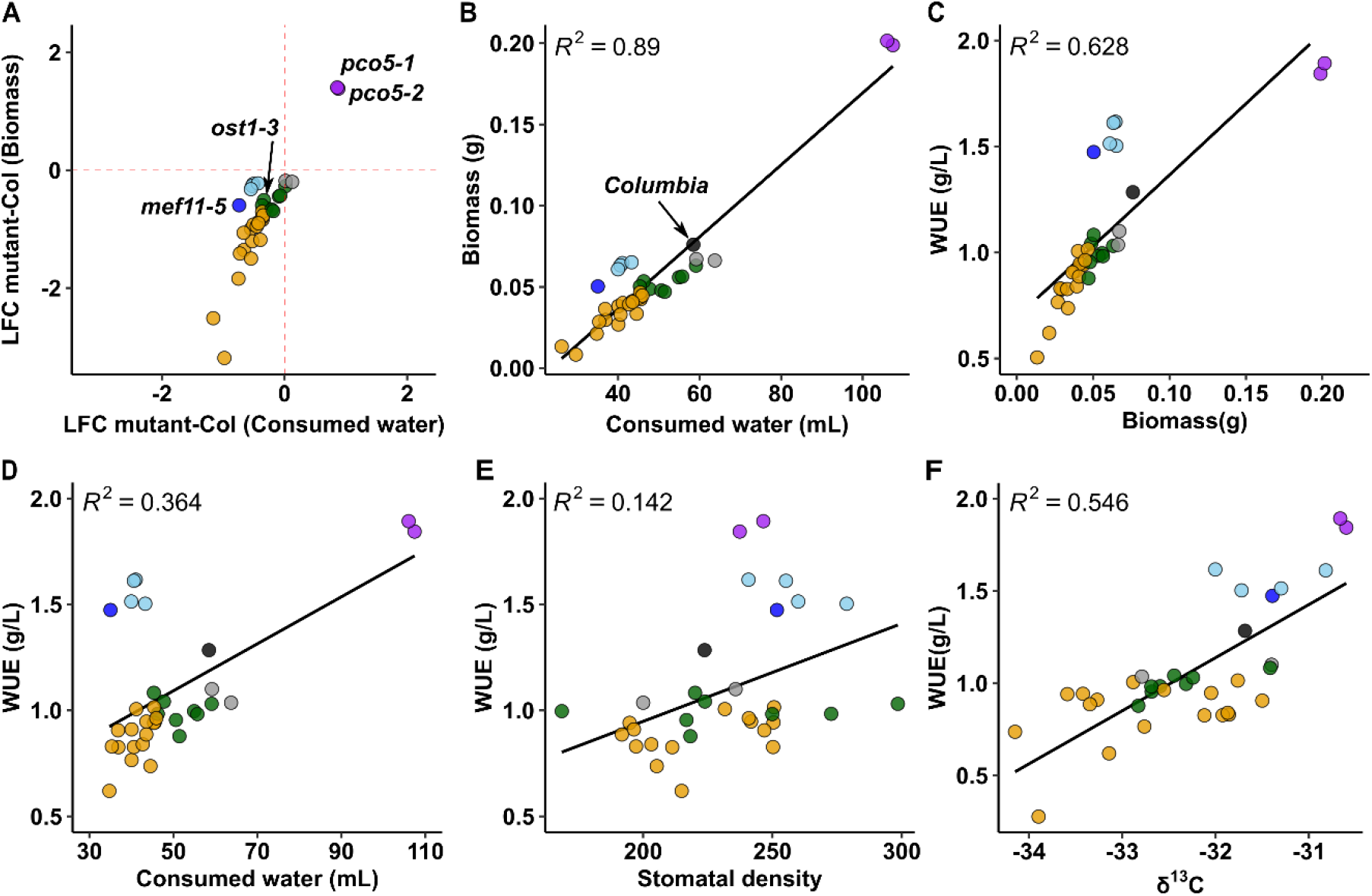
Reverse genetic tests of candidate genes for WUE by gravimetric analysis using larger containers. (A) Relative contribution of biomass and consumed water of a given mutant as a factor leading to altered WUE (g/L, biomass produced per water consumed). Graph shows the Log2 fold changes (LFC) of above ground biomass and consumed water for a given mutant compared to Col-0 reference (dotted red lines). The mean value for Col-0 and individual T-DNA lines of each gene were plotted (*n* =5-10 biological replicates for each data point). The altered WUE, changes in biomass, and consumed water is indicated by color filled circles; high WUE with increased biomass and increased water consumption (purple), high WUE with decreased biomass and reduced water consumption (dark blue), high WUE with no changes in biomass but reduced water consumption (light blue), low WUE with decreased biomass and no change in consumed water (green), low WUE with decreased biomass and reduced water consumption (orange), low WUE with no changes in biomass or water consumed (grey) (B) Association of above ground biomass with consumed water. (C) Association of WUE with above ground biomass and (D) consumed water. (E) Association of WUE with stomatal density. Note that, for a few mutants with reduced growth, the number of samples is less (*n*=3-4 biological replicates from two independent experiments) because of difficulty in obtaining epidermal peels from those mutants. Three other mutants were omitted from this analysis for the same reason (Fig. S6). (F) Association of 13C isotope composition (δ^13^C) in above ground biomass with WUE (ratio of biomass to water consumed). δ^13^C was measured for the above ground biomass as shown in B. Colors in B to F are as in A except for the Col-0 which is represented by black filled circle.

### Genes that act as negative effectors of WUE

We found that mutants of *Plant cysteine oxidase 5 (pco5-1 and 2), Defective UGE in root (dur-1), Calcium-dependent protein kinase 23 (cpk23)*, and *Nuclear speckle localized RNAa* (*nsra-1 and 2*) had increased WUE, indicating that the proteins encoded by these genes have a negative effect on WUE (Fig. 3). None of these mutants affected SD (Fig. S6A). Two mutant alleles of *PCO5* had a strongly significant increase in gravimetric WUE and δ^13^C while also producing more biomass and consuming more water. (Fig. 2F, purple circles). Thus, mutants of *PCO5* were more water productive despite having significantly reduced leaf temperatures (Fig. 3 and Fig. S7) indicative of a higher transpiration rate. Since *pco5* mutants had no effect on SD, the reduced leaf temperature suggests that stomatal size regulation of stomatal aperture may instead be altered. *PCO5* is one of five cysteine oxidases involved in N-end rule protein degradation and hypoxia response (36).

**Figure. 3.**
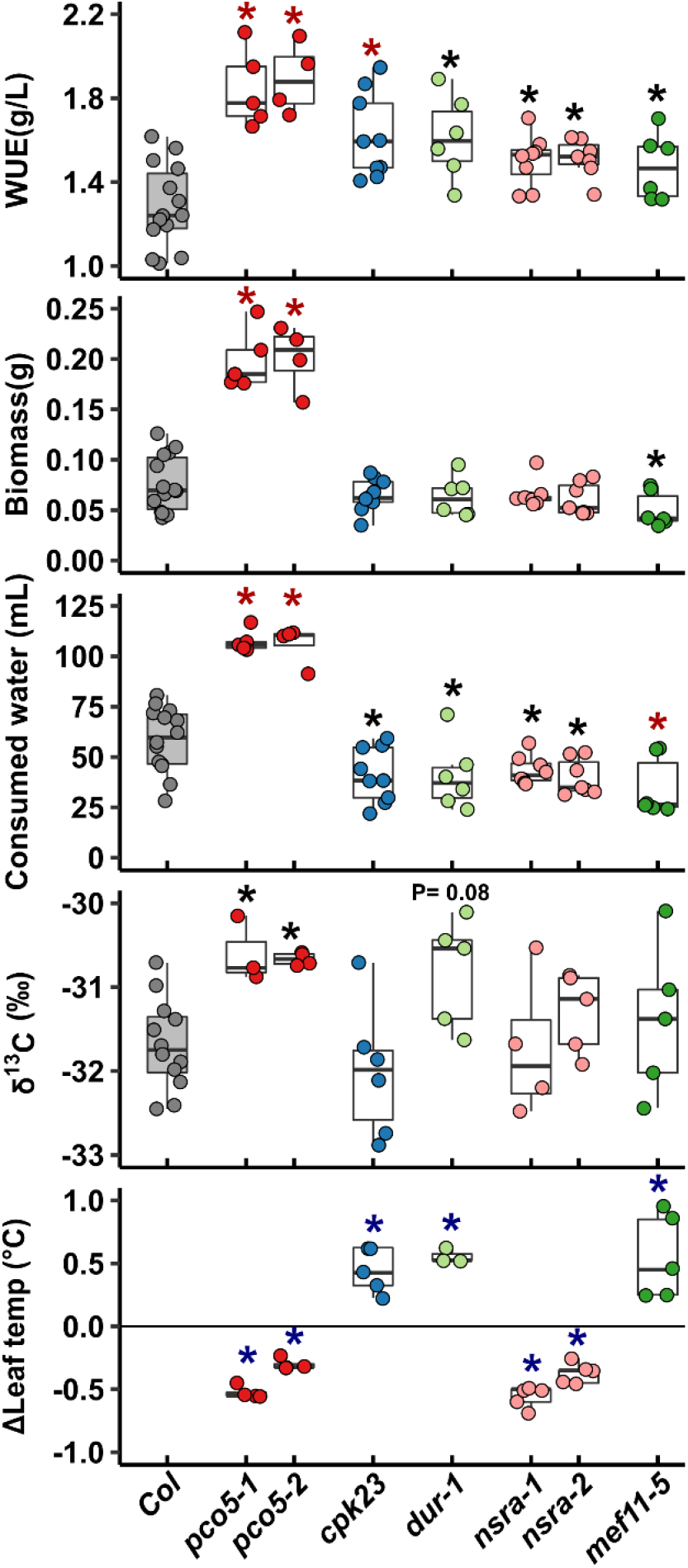
Mutants of four genes had higher WUE than wild type. WUE, above ground biomass, consumed water, δ^13^C analysis for the above ground biomass and leaf temperatures of the plants for which WUE was measured are presented. The box plots indicate the 1^st^ and 3^rd^ quartiles with a median line and the whiskers represents the 1.5x interquartile range. Asterisks indicate statistically significant difference for a comparison of mutants against Col-0 wild type (red: strongly significant difference (adjusted *P<*0.05), black: moderate effect based on nominal *P* value (Dataset S12). P values are listed for the lines with marginally non-significant effect. Note that, the leaf temperatures data was collected from the second independent experiment (*n*=3-5 biological replicates). The difference of leaf temperatures (Δ = mutant – Col) are plotted. The blue asterisks denote a significant difference (*P <* 0.05) based on t-test performed on original leaf temperature values (Plots of original values and thermogram images are provided in Fig. S7)

Mutants of three other genes in this category, *cpk23, dur-1* and both *nsra* alleles, had significantly increased WUE. These mutants had no effect on biomass but showed moderately reduced water consumption suggesting that they altered the transpiration side of WUE (Fig. 3 and Fig. 2F: blue circles). Among these, *cpk23* had the strongest effect on WUE. *dur-1* had a marginally non-significant (*P* = 0.08) increase in δ^13^C consistent with the gravimetric analysis. *cpk23* and *dur-1* had elevated leaf temperatures whereas *nsra-1* and *nsra-2* had decreased leaf temperatures compared to wild type (Fig.3 and Fig. S7). The control mutant *mef11-5* had a moderate decrease in biomass and strongly reduced water consumption leading to increased WUE. It had no effect on SD (Fig. S6A) but did have elevated leaf temperature (Fig. 3 and Fig.S7), consistent with previous reports (32, 33). The elevated leaf temperatures of *cpk23, dur-1* and *mef11-5* are consistent with their reduced water consumption but the decreased leaf temperatures for the *NSRa* mutants contrasts with their reduced water consumption. It is possible that although *NSRa* mutants did not have significantly reduced biomass, they may have had reduced leaf area that could explain the apparent mismatch between water consumption and leaf temperature. *DUR* encodes a UDP glucose epimerase (UGE) involved in UDP-arabinose biosynthesis, possibly affecting cell wall properties (37). NSRa is a RNA binding protein with no previous information to connect it to stress responses or stomatal development. NSRa, and related NSRs, affect alternative splicing and thus could influence activity of downstream genes leading to increased WUE.

### Mutants related to hormone, calcium or stress signaling had decreased growth leading to decreased WUE and water consumption

Mutants of 13 genes had a strong decrease in WUE mainly due to a strong reduction in biomass even though their water consumption was also reduced (Fig.4, and Fig. 2A, orange circles). None of these mutants affected SD (Fig. S6B). Mutants of first five genes in this category *mterf defective in Arabidopsis* (*mda1-3*), *cytochrome P450, family 7070a3* (*cyp707a3-3*), *dehydration responsive element-binding protein 2a* (*dreb2a-1 and 2*), *general control non-repressible 20 (gcn20-2)*, and a hypothetical protein with unknown function (*at1g49170*) had strong decrease in biomass as well as strongly reduced water consumption (Fig. 4). However, biomass was more strongly reduced, leading to substantial decrease in WUE in these mutants.The decreased δ^13^C in these mutants was consistent with the gravimetric WUE. *MDA1, CYP707A3 and DREB2a*, were previously reported to affect ABA and water stress signaling whereas *GCN20* has roles in hormone signaling (38). *MDA1* encodes for mitochondrial transcription termination factor (mTERF). *mda1* was previously shown to enhance salt and osmotic stress tolerance probably due to reduced sensitivity to ABA (39). *CYP707A3* acts as an ABA 8’-hydroxylase involved in ABA catabolism (40). *cyp707a3-3* had elevated leaf temperatures (Fig. 4, Fig. S8) consistent with its reduced water consumption and with previous observation of lower stomatal conductance and higher basal ABA levels in unstressed *cyp707a3* plants (40). *DREB2a* is a transcription factor that functions in both water and heat-stress responses (41). Of the two *dreb2a* alleles we examined, *dreb2a-3* had stronger effect than *dreb2a-4* consistent with *dreb2a-4* being a knockdown, rather than knockout (42). Both mutant alleles of *DREB2a* had decreased leaf temperatures (Fig. 4 and Fig. S8), indicating that they may affect WUE by controlling stomatal aperture. *GCN20* encodes an ABC transporter family protein. Mutants of *GCN20* were defective in MAMP/bacterium-triggered stomatal closure but respond normally to salicylic acid (SA) and ABA (38). *gcn20-2* had decrease in leaf temperatures and no change in SD, consistent with previous report (38). The *AT1G49170* mutant also had decreased leaf temperature (Fig. 4, Fig. S8).

**Figure. 4.**
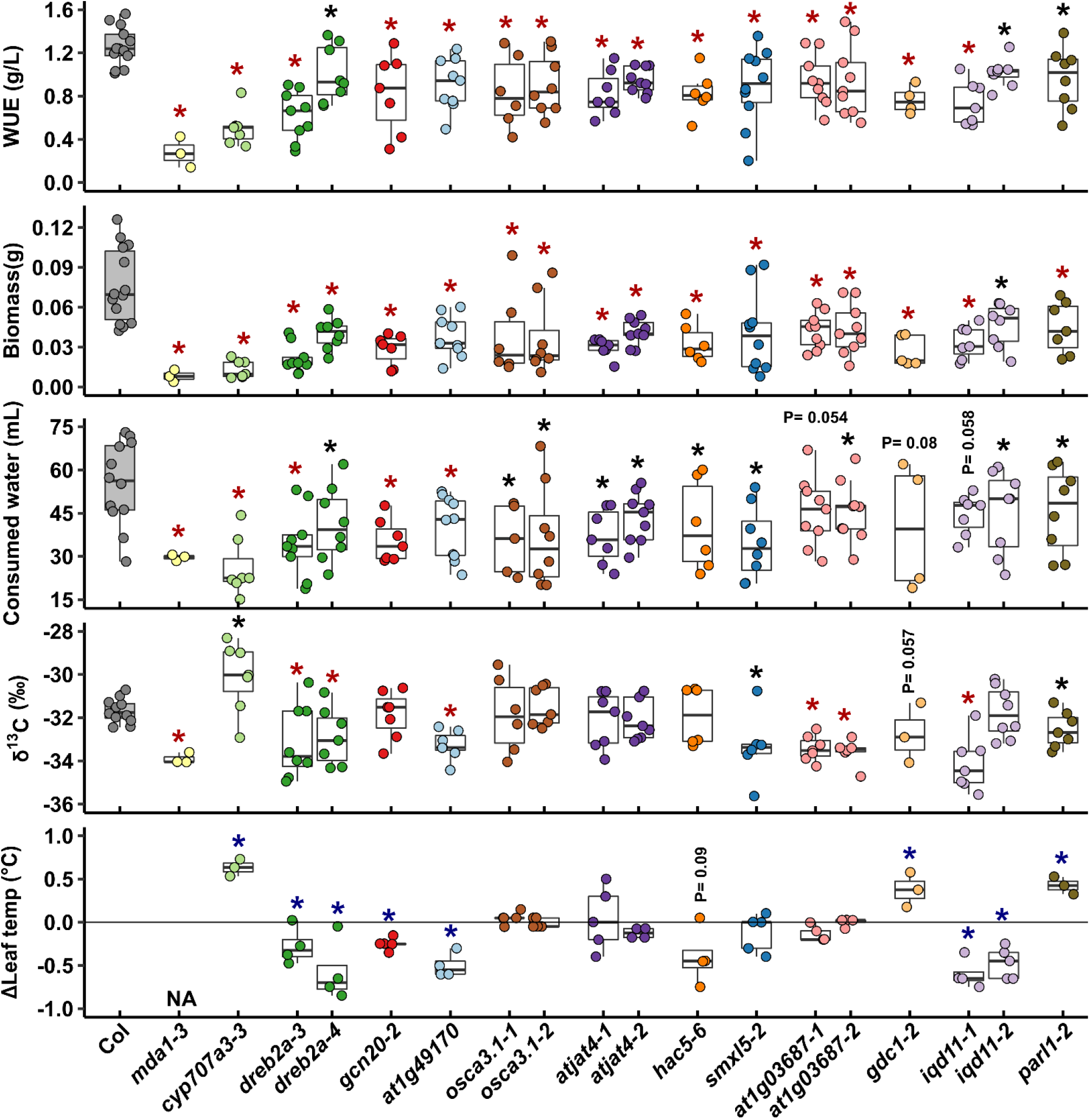
Mutants of thirteen genes had reduced biomass and reduced water consumption as a factor leading to decreased WUE. The data presentation format is the same as described for Fig. 2. Plots for original leaf temperature values and thermogram images are provided in Fig. S8 and S9. NA denotes missing data (“Not available”).

Mutants of the next six genes in this category include *osmosensitive calcium-permeable cation channel 3*.*1/early responsive to dehydration 4* (*osca3*.*1*/*erd4*), *jasmonic acid* (JA) *transporter 4* (*jat4/atjat4*), *histone acetyl transferase* (*hac5*), *suppressor of max2 1 – like 5* (*smxl5*), *AT1G03687*, and *grana deficient chloroplast 1* (*gdc1*) had strong decrease in biomass with moderately reduced water consumption, indicating that biomass was the driver for the decreased WUE of these mutants (Fig. 2A, orange circles). OSCA3.1 and JAT4 have known or highly probable roles in stress or hormone signaling (43, 44) and the mutants of *OSCA3*.*1* and *JAT4* showed strong decrease in WUE. Similarly, *hac5-6, smxl5-2*, mutants of *AT1G03687*, and *gdc1-2* all had strongly decreased WUE. HAC5 functions in the transcriptional repression of genes related to flowering time and floral development (45). *hac5-6* had marginally non-significant decrease in leaf temperatures (*P* = 0.09) (Fig. 4, Fig. S9). SMXL5 promotes secondary phloem formation (46) and *smxl5-2* had decreased δ^13^C consistent with gravimetric WUE. The altered WUE in *smxl5-2* might be due to upregulation of several stress related pathways in this mutant (46). The function for AT1G03687 is unknown. Both mutants of *AT1G03687* had strong decrease in δ^13^C consistent with gravimetric WUE. GDC1 is a ankyrin domain containing chloroplast protein essential for grana formation (47). *gdc1-2* had marginally non-significant decrease in water consumption (*P* = 0.08) and δ^13^C (*P* = 0.057) and showed increased leaf temperature. Consistent with previous reports, *gdc1-2* exhibited pale green leaves, and ceased growth at the vegetative stage (47), indicating that this mutant had decreased WUE because of impaired photosynthesis.

The next gene in this category was *IQ-domain 11* (*IQD11*), another Ca^2+^ responsive protein involved in regulation of microtubule orientation (48). Among two T-DNA alleles, only *iqd11-1* showed strong decrease in WUE, biomass and δ^13^C, while *iqd11-2* had a more moderate effect on these phenotypes and was not significant for δ^13^C. *iqd11-2* has a T-DNA insertion in the 5’ untranslated region (UTR) and thus is likely a knockdown rather than knockout mutant (Dataset S11). Note that, *iqd11-1* had marginally non-significant decrease in water consumption (*P* = 0.058). Both mutants of *IQD11* had decreased leaf temperatures (Fig. 4, Fig. S9). *Parallel 1/Nucleoline-1 (PARL1/NUC1)* is involved many aspects of ribosomal biogenesis and affects leaf venation (49). *parl1-2* is the only mutant in this category that had a moderate decrease in WUE. It had decreased δ^13^C consistent with gravimetric WUE and showed elevated leaf temperature (Fig. 4, Fig. S9).

The changes in leaf temperatures for mutants in this category suggests their role in stomatal response ((Fig. 4, Fig. S8 and S9). The elevated leaf temperatures found for *cyp707a3-3, gdc1-2* and *parl1-2* was consistent with their reduced water consumption; however, decreased leaf temperatures found for mutants of *DREB2a, GCN20, HAC5* and *IQD11* contrasts with their reduced water consumption. Here again, change in leaf area or rosette architecture may explain the seeming mismatch between leaf temperature and water consumption.

### Mutants with reduced WUE and biomass but with little effect on total water consumption

Mutants for six genes, *With No Lysine* (*WNK11*), *Plethora 1* (*PLT1*), *AT2G45460, AT3G62220, Basic Penta Cysteine* (*BPC2*) and *Carbon Catabolite Repressor 4b* (*CCR4b*), had decreased WUE and reduced biomass but no significant change in water consumption (Fig.5 and Fig. 2A: green circles) or SD. WNK11 belongs to WNK protein kinase family that are involved in various processes, such as ABA sensitivity, proline accumulation, regulation of flowering time and intracellular signalling through G protein (50); however, the function of WNK11 itself is unknown. *wnk11-1* had strongly decreased WUE and biomass while others had moderate effects in this category. The decreased δ^13^C in *wnk11-1* was consistent with gravimetric analysis. PLT1 is a negative regulator in ABA inhibition of root growth (51). *plt1* had marginally non-significant (*P* = 0.054) decrease in δ^13^C consistent with gravimetric WUE. BPC2 functions in negative regulation of osmotic stress tolerance and it also determines β-1,4-galactan accumulation in response to salt stress (52, 53). *bpc2* also had decreased δ^13^C consistent with gravimetric WUE. It is possible that WNK11, PLT1 and BPC2 affect WUE through mechanisms related to ABA or osmotic stress signalling.

*AT2G45460* and *AT3G62220* encode proteins of unknown function. For *AT3G62220*, we observed disparate results for two independent T-DNA lines; only *at3g62220-*1 (promoter insertion) had a significant effect on biomass, while *at3g62220-2* (exonic insertion) had marginally non-significant (P = 0.08) effect. Conversely, *at3g62220-2* had significant effect on leaf temperature and SD while *at3g62220-1* had no significant effect (Fig 5, Fig. S6C and S10). *ccr4b-1* had marginally non-significant decrease in WUE (*P* = 0.08) and it had decreased leaf temperatures. CCR4b determines the poly (A) length of transcripts related to starch metabolism and may also affect degradation of stress responsive RNAs via its interaction with Pumilio RNA-binding protein 5 (54). The control mutant *ost1-3* had moderate effect on WUE and biomass, strong decrease in δ^13^C and expected decrease in leaf temperatures (Fig. 5 and Fig S10). Surprisingly the altered leaf temperatures did not affect the overall water consumption in these mutants and none of these mutants influenced SD except for *at3g62220-2* that had increased SD (Fig.S6C). There is no prior information for genes in this category to link them to WUE or traits related to plant water relations.

**Figure. 5.**
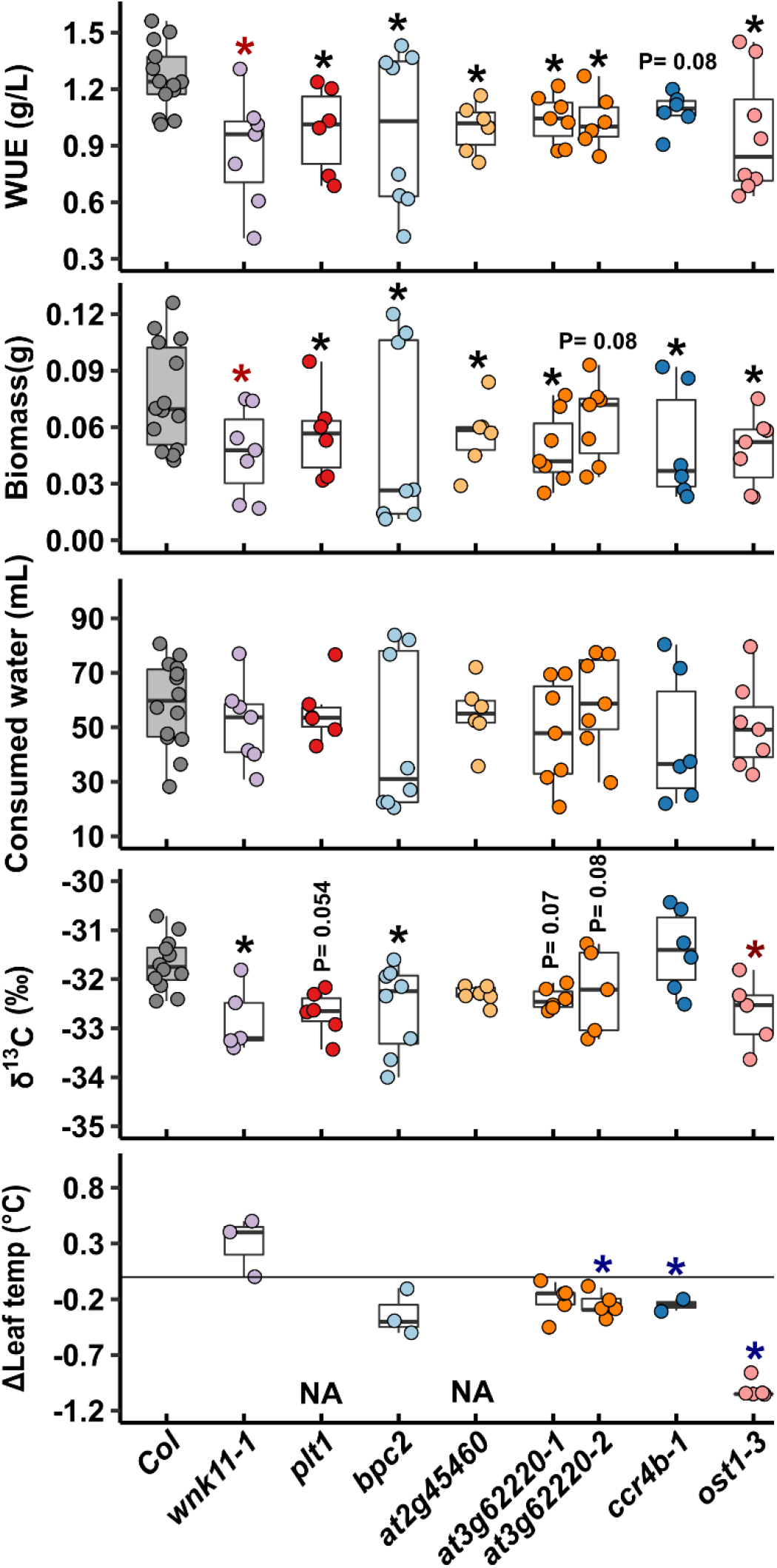
Mutants of six genes that had reduced biomass as a factor leading to decreased WUE. The data presentation format is the same as described for Fig. 2. Plots for original leaf temperature values and thermogram images are provided in Fig. S10. NA denotes missing data (“Not available”).

Finally, we found mutants of Natural resistance associated macrophage protein 4 (NRAMP4) and Carbamoylphosphate synthetase subunit B/Venosa3 (*CarB/VEN3*) had marginally non-significant decreases in WUE without significantly affecting final biomass or water consumption (Fig. S11 and Fig. 2A: grey circles). In addition, these mutants did not affect δ^13^C or SD (Fig. S11 and S6D) and *carb* did not affect δ^13^C or leaf temperatures (Fig. S11 and S11). NRAMP4 is functionally redundant with NRAMP3 in maintaining photosynthesis under cadmium and oxidative stress (55). CarB is involved in the conversion of ornithine into citrulline in arginine biosynthesis, however *carB* mutant used in this study is a knockdown mutant that expresses reduced level of carB (56) suggesting that a *carB* knockout mutant may have greater effect on WUE.

## Discussion

Our strategy of combining GWAS results and gene expression generated a broader set of candidates for reverse genetic tests. We found 25 genes with significant differences in WUE (Dataset S9 and S11). The success rate is moderate [positive rate of 0.35 (25/70 discovered candidates)]. However, the lack of phenotype in T-DNA mutants cannot conclusively rule out a candidate gene as the source of the GWAS association since knockout mutants might not recapitulate gain-of-function or altered function natural alleles. Also, gene redundancy could be another mitigating factor making the impact of knockouts of some of our candidate genes. Construction of higher order mutants including close homologs or the generation of overexpression lines might reveal phenotype for these genes where T-DNA mutants did not alter WUE. Despite this caveat, a combined GWAS and reverse genetic approach proved to be an effective way to discover new genes influencing WUE without relying on assumptions about underlying mechanisms, as it has been for other stress-related traits (23, 24). The candidate genes we validated belong to a range of gene families such as histone acetyl transferases, mitochondrial transcription factors, mechanosensitive ion channels, ABC transporter G family, ankyrin repeat-domain containing proteins, cytochrome P450, family 707A. Strikingly, a recent GWAS study also identified genes belongs to these families as some of the most promising candidates for WUE related traits in sorghum (25).

Since WUE is a ratio of biomass accumulated per unit of water consumed, it can be influenced by factors that alter either side of the ratio (or affect one side of the ratio more than the other side). Our approach allowed us to find mutants affecting either side of the WUE ratio and to parse out which aspect of WUE was most affected by each mutant. Interestingly, we found mutants affecting both sides of the WUE ratio, thus demonstrating that changes in growth are as likely to influence WUE as changes in transpiration. In addition, we did not find any mutants where decreased biomass was associated with increased WUE. Thus, contrary to what may be a common assumption, our data shows that simply decreasing biomass was not sufficient to increase WUE. This pattern also supports the hypothesis that our approach identified specific effectors of WUE, as opposed to merely finding plants that cannot grow well. Our data identified more mutants with decreased WUE than with increased WUE, suggesting that WUE is under positive regulation as part of mechanisms balancing net carbon assimilation to water use. None of the genes we identified would likely have been predicted to affect WUE *a priori* (with possible exception of *CYP707a3*).

While the new WUE effector genes we identified may at first look seem to be a random assortment of genes, they can in fact be grouped into several categories that make their potential roles in WUE clearer. The first group of genes we identified are genes involved, or potentially involved, in stress-related signaling or hormone metabolism but not previously connected to WUE. The most prominent of these is *PCO5*. The *pco5-1* and *pco5-2* mutants had the strongest increase of WUE of any mutants we examined because of their strongly increased biomass production. PCOs are components of the N-end rule pathway which act as redox status sensors. Several PCOs act as oxygen or redox status sensors that regulate stability of the Group VII ethylene response factors (ERF-VII) via N –end rule pathway of targeted proteolysis in response to hypoxia (57). PCO5 activity towards ERF-VII destabilization was demonstrated *in vitro*, but it’s *in planta* function remains to be established (58). Interestingly, Proteolysis 6 (PRT6) and other N-end rule components were shown to affect ABA accumulation and ABA sensitivity (24, 59). Thus, our data provide another piece of evidence that the N-end rule pathway influences many types of environmental responses in addition to the hypoxia responses where it has been best characterized. Whether or not the increased biomass and WUE of *pco5* mutants is dependent on ERF-VII protein stability or other targets of N-end rule degradation will be of interest for future studies.

Our data also implicates calcium-dependent signaling in WUE. Our finding that OSCA3.1 had strongly reduced WUE and biomass is especially interesting as there has so far been little physiological data to link the short-term calcium responses mediated by OSCAs to longer-term acclimation to water limitation or other environmental conditions. Also, mutants of the calcium responsive protein IQD11 had similar effect on WUE and biomass as *osca1*.*3* mutants. As most IQDs are also microtubule binding proteins, IQD11 could also affect WUE via effects on microtubule stability or organization that alter stomatal function or affect biomass (60, 61). Mutation in one of the calcium dependent protein kinases, *cpk23* led to increased WUE in our growth conditions. It is surprising that *cpk23* had increased WUE rather than decreased WUE as seen in *ost1* (*snrk2*.*6*) because CPKs and SnRK2 can both activate the guard cell slow anion channel-associated 1(SLAC1) by phosphorylating distinct sites on SLAC1 (62). While this suggests a similar function of CPK23 and SnRK2s, our results are consistent with another study which found that *cpk23* had slower water loss and reduced stomatal aperture (63) Together, these findings that OSCA3.1, IQD11 and CPK23 all strongly affected WUE, but affected it in different ways, suggests that multiple calcium-dependent changes in guard cell function or regulation of growth (biomass) can alter WUE. Consistent with this idea, the genes associated with the top 500 GWAS SNPs also included CPK8 and several other types of calcium binding proteins (Dataset S2).

Other signaling genes we found to impact WUE include several genes that may have been expected to have some effect on WUE. Nevertheless, the actual mutant phenotypes we found were still surprising in several ways. CYP707A3 is involved in ABA catabolism and the simplest expectation may be that even a slight increase of endogenous ABA levels of *cyp707a3* would enhance stomatal closure and increase WUE. We saw no evidence of this but instead observed that *cyp707a3* had decreased WUE driven by a strong decrease in biomass accumulation. This suggests that the biomass inhibition caused by disrupted ABA metabolism in *cyp707a3* was too substantial to be overcome by any increase of stomatal closure. Similarly, *PLT1* mutant had enhanced ABA inhibition of root growth but *plt1* had decreased WUE and biomass similar to *cyp707a3*. The WNK kinase family is known to affect ABA sensitivity but there is no specific information on WNK11 (50). JAT4 is involved in JA translocation and there are a number of indications that JA affects ABA sensitivity (and vice versa) and that the ratio of endogenous JA to ABA accumulation is important for various stress response.

While several of the genes mentioned above have a connection to ABA signaling, it is perhaps surprising that we did not find more ABA-related genes among our GWAS candidates. Moreover, the only mutant that we observed to have decreased biomass and increased WUE was the ABA-hypersensitive mutant *meff11-5* which was used as a control. This suggests that knockout mutants where stomata remain sufficiently closed to limit photosynthesis are relatively rare compared to other loci with more moderate, or indirect effects on WUE. It is also possible that the core ABA signaling components that have strong effects on ABA sensitivity have relatively little natural variation among the accessions used in our GWAS analysis and thus were not detected in our list of top candidate genes. Similarly, another type of gene which may be expected to affect WUE is developmental regulators that determine stomatal density or size. Surprisingly, our GWAS analysis did not find any known regulators of stomatal development that show strong association with δ^13^C. The only known stomatal development genes among the candidate genes identified by GWAS was *SCREAM1/ICE1* (associated with the 21^st^ ranked SNP) and *CKB1* associated with the 59^th^ ranked SNP (Dataset S2). However, these were not analyzed further as they were not among the stress up-regulated genes in the two transcriptome data sets we used to select stress up-regulated candidate genes. Other stomatal regulators may have limited effect on the WUE plasticity that was the basis of our GWAS or may have limited natural variation in the group of accessions studied. The mutants we did find to affect WUE generally had no effect on stomatal density (Dataset S6), further indicating that, at least in the population of Arabidopsis accessions used for our GWAS, stomatal development was not a key driver of WUE variation. An alternative explanation would be that key genes involved in hormone responses or core components of cell fate and development are under strong purifying selection due to potential pleiotropic costs (64). It may be that there is more standing genetic variation in more specialized or modular aspects of physiology or growth that lead to the observed natural genetic variation in WUE

Maximizing the amount of biomass, or other components of yield, produced per amount of water input can be a beneficial agronomic trait in many environments. However, increasing WUE at the expense of biomass production may not be as useful. Even by focusing solely on reverse genetic, loss of function analysis, we could find numerous new genes that maybe useful for more targeted study and manipulation of WUE without negative effects on biomass accumulation. For example, *PCO5* mutants have strongly increased biomass accumulation leading to dramatic increase in WUE. Conversely, we found several mutants that decrease WUE without affecting biomass. These would be good candidates for gain of function analysis. In any case, these results uncover several new pathways that may be used to influence WUE.

## Materials and Methods

### GWAS-mapping

GWAS was performed using the plasticity of WUE (plasticity = drought – well watered control) as well as the WUE of the well-watered control and drought treatments. The data set of WUE (δ^13^C) covering 185 accessions (Dataset S1) was linked to published genomic data on accession from 250K SNP chip and were used in association analyses. These data were previously analyzed at the phenotypic level in Kenney et al. (1). We implemented a genome-wide association study for WUE (δ^13^C) using a linear mixed-model approach (1). The model included random effects for each observation that were constrained to a correlation structure by the genome-wide kinship matrix (identity-in-state). We included SNPs with a minor allele frequency of at least 0.1.

### Water use efficiency (δ^13^C) estimation

The details for the growth condition and experimental design can be found in Kenney et al. (2). In brief, natural accessions of Arabidopsis were grown in green house conditions (16h light/ 8h dark, 1000 *µ*mol m^−2^ S^−1^, 18-21°C) in the cone containers (164 ml soil volume capacity). Plants were treated identically until week 4 before altering the water regime. Then, watering was continued for well-watered wet treatment plants and ceased for drought treatment until week 6. At the end of the experiment, most plants had completed flowering and many were senescing. The above ground material of the entire shoot from available replicates from each accessions was pooled and course ground in centrifuge tubes. After that, subsamples were fine ground in microcentrifuge tubes. Two mg of finely ground tissue was loaded into a tin capsule and analyzed at the UC Davis Stable Isotope Facility (http://stableisotopefacility.ucdavis.edu). Data are presented as carbon isotope ratios relative to the V-PDB standard (*R*_PDB_), where δ^13^C (‰) = (*R*_sample_/*R*_PDB_-1)*1000. These values are expressed per mil (‰).

We followed the same procedure as above to estimate δ^13^C for T-DNA mutants from the large container experiments in the current study except that we estimated δ^13^C for each available replicates of T-DNA mutants instead of pooling them together as it was done in Kenney et al. (1) for natural accessions. See details below for experimental design and growth conditions.

### Prioritization of GWAS candidates and enrichment analysis

Prioritization of GWAS candidates for the functional validation is explained in details in the result section. In brief, we ranked the genome for candidate genes impacting WUE plasticity by sorting the SNPs by nominal p-value. We focused on the genomic regions tagged by the top 500 SNPs and gene space in tight LD (corresponding to an LD window of 5 Kb on either side of tag SNPs). This effort identified 1058 SNPs. Using existing drought stress transcriptome data (2,3), we identified 198 genes in these genomic intervals that were drought upregulated (Dataset S5). These 198 genes were further subjected to Gene Ontology (GO) enrichment using DAVID (Database for Annotation, visualization, and Integrated Discovery) bioinformatics resources v6.8 with high classification stringency (3, 4) to identify the most relevant biological terms of this candidate set. We retrieved 10 significant functional annotation terms listed according to their enrichment (adjusted *P <* 0.05) (Fig. S2C and S3).

### T-DNA screening

T-DNA insertion lines were obtained from the Arabidopsis Biological Resource Center (ABRC). We sought to obtain multiple T-DNA lines for each candidate gene and further prioritized the lines with a T-DNA insertion in the exonic region of a gene to increase the possibility of retrieving knockout mutants. However, obtaining multiple lines and lines with exonic insertion was not always possible due to lack of suitable T-DNA mutants or inability to isolate homozygous individuals (Dataset S8). In total, we prioritized 101 genes for functional follow up (See the main text for details). Of these selected genes, T-DNA mutants for 15 genes were found to be associated with multiple loci or ABRC stocks were not available (Dataset S8, red highlighted). So, we screened a total of 152 T-DNA lines of 85 genes (Dataset S8) and identified 88 homozygous lines covering 70 candidate genes. We used CELLSTAR® 48 well cell culture plates (Greiner Bio-one, North America, Inc) for screening homozygous T-DNA lines. The plates were prepared by pipetting 0.5 ml of autoclaved half-strength Murashige-Skoog (half-MS) medium to each well. We planted one seed per well, and 24 seeds were initially used per T-DNA lines for genotyping. After planting seeds, the plates were sealed using MicrosporeTM Surgical Tape; 3M Health Care, USA) to hold the plate lid and the base, and the entire setup was cold stratified for four days in a 4°C refrigerator. Plates were then moved to the growth chamber (16h light period, 23°C, light intensity of 100–120 *μ*mol m^−2^ s^−1^)). After 2 weeks or when plants were at the 4-leaf stage, one small leaf was collected in 1.5 ml tube containing 200 μL of Edward’s buffer (200 mM Tris HCl pH 7.5, 250 mM NaCl, 25 mM EDTA, 0.5% SDS). The tissue was macerated using disposable polypropylene pestles to obtain a pale green-colored homogenate.

Further, 1 μL of this homogenate was included in PCR analysis. We followed the PCR genotyping of T-DNA insertion as described in O’Malley et al. (5), and primers for genotyping were generated by the SIGnAl web resource; http://signal.salk.edu/tdnaprimers. Homozygous lines were then transferred from plates to the soil. Plants were grown to maturity for seed harvesting. To homogenize the age of seeds, we again grew all homozygous lines at once to harvest seeds for phenotypic analysis.

### WUE measurement by gravimetric analysis

We isolated 88 homozygous T-DNA insertion lines covering 70 candidate genes (Dataset S9 and S10). Water use efficiency for all homozygous T-DNA lines was determined by gravimetric analysis. To accommodate all replicates in two growth chambers, we initially grew plants in 50 ml falcon tubes as a small container experimental system (Fig. S4B). This system was effectively used to determine water use efficiency in Arabidopsis (6, 7), and we adapted the system with few modifications. In brief, a standard potting mix was combined with 20% Turface and soil was well moistened to grow plants for six weeks without further supply of water. We used a ratio of soil (8): Turface (2): water (6) to prepare an evenly mixed, well-moistened soil mixture. Tubes were filled with the soil mixture to the brim. Tubes were then capped and wrapped with aluminum foil to prevent algal growth. Next, 4-6 seeds were placed in a hole (approximately 2 mm wide) made in the cap by drilling. The tubes were randomized and placed in racks (25tubes/rack). The racks were then placed in a closed transparent container with a water-covered bottom. The entire setup was moved to a walk-in cold chamber for cold stratification of the seeds. After 4 days, racks were moved to a short day growth chamber (8h light period, 23°C, light intensity of 100–120 *μ*mol m^−2^ s^−1^). Racks were cycled within and between the chambers every two days to minimize the impact of microenvironment variation. Excess seedlings were thinned on the 7th day to leave one seedling per tube, and tube weight was measured and recorded as W0. After 4 weeks, rosettes were decapitated and dried for 3 days at 65 °C oven. After removing the rosette, tube weight was recorded as W. Water loss was calculated as W0 (g) – W(g). For water, 1 g is equal to 1 ml. Water use efficiency was calculated as a dry aboveground biomass (mg) over ml of water used (biomass/water consumed).

After the initial screening, T-DNA lines differing in WUE compared to were tested for WUE in large plastic cups (237 ml soil volume) as a large container experimental system (Fig. S4C), more details are given in the results section). Our goal was to provide more soil nutrients and a larger growing surface and test whether the WUE phenotype previously observed was stable to growth conditions. Larger containers with lids also allowed us to easily use thermal imaging to capture rosette temperatures by avoiding complications caused by soil background. We performed two independent experiments with 5 biological replicates per genotype. The methods and growth conditions were largely similar to the small container system with a few modifications. In brief, seeds were incubated at 4 °C in 1.5 ml tubes containing 1 ml of water. After 4 days. A 237 ml plastic cup was filled with a soil mixture (soil (8): turface (2): water(6), and 5-10 seeds were planted in the hole made to center of the lid using a forged steel hollow punch (W.W. Grainger, Inc, USA). The containers were randomly distributed in the growth chamber and shuffled within and between the chambers every two days. Growth condition and duration of the experiment was similar to the small container system. At the end of the experiment, we sampled one randomly selected fully grown leaf for stomatal density measurements before decapitating the rosette for biomass weight. We collected thermal imagery of rosettes before harvesting only for the 2^nd^ independent experiment, before collecting samples for stomatal density and biomass measurements. Note that, we dried rosettes for 4 days at 65 °C rather than 3 days as mentioned for the small container system. WUE was calculated as mentioned above.

### Stomatal density and stomatal index measurements

Images to evaluate the number and size of stomata and mesophyll cells were captured with a Nikon Eclipse Ni microscope equipped with a Nikon DS-Ri2 color camera at 20x magnification. One randomly selected fully grown leaf of the same developmental stage was sampled to prepare each slide for epidermis phenotyping. Clear nail polish was applied to the abaxial surface of leaf and allowed to dry for 10 minutes at room temperature. The dried nail polish area was peeled off with clear tape to prepare slides for microscopy. Images were analyzed with ImageJ software to measure cell length and width in µm, which were then used to calculate stomatal and epidermal cell area. We measured one randomly selected stomata and epidermal cell from each leaf. Stomatal area and epidermal cell area were calculated using the following equation:

Cell Area (µm2) = Cell Length x Cell Width. We then counted every stomata and epidermal cells observed under the microscope field and also measured the area of the total view of the microscope field. We calculated total stomatal cell and epidermal cell area by multiplying the total number of cells under a microscopic field by the single cell area. The ratio between stomatal to epidermal cell area was calculated by dividing total stomatal area by total epidermal cell area. The following measurements were used to calculate other leaf anatomy traits:

Stomatal density = number of stomata in entire FOV / area of total microscope field (µm2) Stomatal Index (%) = Stomatal density *100/Stomatal density + Epidermal cell density”

### Thermal imaging and leaf temperatures measurement

Plants from the second large container experiments were used for thermal imaging and leaf temperatures measurement. For imaging, biological replicates of a mutant were gathered with Col-0 wild type to comprise a single field of view of the camera and images were taken using a FLIRA325sc thermal camera. Leaf temperature was measured using FLIR ResearchIR Max software. We used the freehand ROI (Region of Interest) tool to trace the entire rosette carefully to avoid background and obtained the mean temperature for individual rosettes.

### Statistical analysis

WUE data of T-DNA mutants from the small container experiment was analyzed by fitting a linear mixed model using the “lmer” function in the “lme4” package in R (8). We included T-DNA insertion lines nested in gene as fixed effect and used container racks as a random effect. Estimated marginal means (EMMs, also known as least-squares means) were derived from the result of the mixed model and significance of T-DNA insertion line was tested using planned contrasts on each gene level versus Col-0 wild type using R package “emmeans” (https://github.com/rvlenth/emmeans). The Type-1 error rate (alpha=0.05) for multiple testing was controlled using a strict Bonferroni correction (Dataset S9). Results for the data analysis at the T-DNA level (EMMs and statistical significance for each T-DNA lines), where T-DNA was used as fixed effect is given in Dataset S10.

For the phenotypic measurements (WUE, biomass, water consumed, *δ*^13^C, Stomatal density), from the large container system, we used a similar analysis approach as above. We simply used individual T-DNA as a fixed effect and experiment cohort as a random effect, and the pairwise comparison with Col-0 wild type as reference was conducted at the T-DNA level (Dataset S12)

Linear regression was performed in R using “lm’ function for all the correlation analysis. T-test for leaf temperatures data of each mutant against Columbia wild type was performed in R using “compare means” function.

## Supplemental data files

Figures S1 to S12

Datasets S1 to S12

## Author Contributions

G.B.B., and T.E.J., designed research; G.B.B., S.R., L.Z., T.H., G. Z.C., and J.E.B. performed research. G.B.B., J.R.L., S.R., and T.H. analyzed data. G.B.B., P.E.V., and J. wrote the paper, with input from J.R.L. T.H., and S.R. contributed to writing.

## Acknowledgments

We thank Dr. Annie Marion-Poll (Institut Jean-Pierre Bourgin, INRA), for *mef11* seeds; Dr. Shuhua Yang (China Agricultural University) for *ost1-3* seeds. We also thank Dr. Ying-Jiun C. Chen and all our lab members for critical reading and constructive comments on the manuscript. This work was supported by funding from the National Science Foundation (DEB-0420111) and (IOS-0922457) to T.E.J.

